# Elevation of intracellular levels of nitric oxide in SHR attenuates hyperproliferation of vascular smooth muscle cells through the inhibition of AT1 receptor expression and c-Src/growth factor receptor signaling pathways

**DOI:** 10.1101/474163

**Authors:** Ekhtear Hossain, Oli Sarkar, Yuan Li, Madhu B. Anand-Srivastava

**Affiliations:** Department of Pharmacology and Physiology, Faculty of Medicine, University of Montreal, Montreal, Canada.

**Keywords:** SNP, VSMC, hyperproliferation, Cell cycle proteins, AT1 receptor, SHR.

## Abstract

We previously showed that decreased levels of intracellular nitric oxide (NO) contribute to the hyperproliferation of vascular smooth muscle cells (VSMC) from spontaneously hypertensive rats (SHR). The present study investigates if elevation of intracellular levels of NO by *in vivo* treatment of SHR with NO donor, sodium nitroprusside (SNP) that was shown to attenuate hypertension could attenuate the hyperproliferation of VSMC and identify the molecular mechanisms. Intraperitoneal injection of SNP (0.5 mg/kg BW) into 8-week-old SHR and WKY rats twice a week for two weeks increased significantly the intracellular levels of NO in aortic VSMC and resulted in the attenuation of hyperproliferation of VSMC from SHR to control levels. The antiproliferative effect of SNP was associated with the restoration of the overexpression of cell cycle proteins, cyclins D1, E, Cdk2, Cdk4, phosphorylated pRB and decreased expression of Cdk inhibitors p21^Cip1^ and p27^Kip1^ towards control levels. In addition, SNP treatment also attenuated the overexpression of angiotensin II receptor type 1 (AT1) receptor, phosphorylation of c-Src, EGF-R, PDGF-R, IGF-IR and ERK1/2 in VSMC from SHR to control levels. These results suggest that the augmentation of intracellular levels of NO elicits antiproliferative effect that may be mediated through its ability to inhibit the enhanced expression of AT1 receptor, activation of c-Src, growth factor receptors and MAP kinase signaling and overexpression of cell cycle proteins.

## Introduction

Vascular remodeling due to the exaggerated proliferation and hypertrophy of vascular smooth muscle cells (VSMC) occurs in several vascular disease states including atherosclerosis, hypertension, and diabetes (1-3). VSMC from spontaneously hypertensive rats (SHR) have been shown to exhibit enhanced proliferation compared with age-matched Wistar Kyoto (WKY) rats (4, 5). The augmented levels of endogenous vasoactive peptides including angiotensin II (Ang II) and endothelin-1 (ET-1) were shown to contribute to hyperproliferation of VSMC from SHR through oxidative stress, transactivation of epidermal growth factor receptor (EGF-R) and mitogen-activated (MAP) kinase signaling pathways (5-7). In addition, the enhanced expression of Giα proteins was also implicated in hyperproliferation of VSMC from SHR because inactivation of Giα proteins by pertussis toxin as well as knockdown of Giα proteins by siRNA attenuated the hyperproliferation of VSMC from SHR (5).

Excessive entrance of cells from G_0_/G_1_ phase to S-phase of cell cycle is associated with hyperproliferation of VSMC from SHR (8). In addition, the cell cycle proteins from G_1_-phase were reported to be overexpressed in VSMC from SHR and implicated in the hyperproliferation (9, 10). We previously demonstrated that the overexpression of Giα proteins and enhanced MAP/PI3 kinase activation contribute to the enhanced proliferation and expression of cell cycle proteins in VSMC from SHR, because PD98059, wortmannin and pertussis toxin, the inhibitors of MAP kinase, PI3 kinase and Giα proteins respectively, attenuated the hyperproliferation of VSMC from SHR and overexpression of cell cycle proteins to control levels (10). In addition, we also showed the implication of oxidative stress, c-Src and growth factor receptor activation in the overexpression of cell cycle proteins in VSMC from SHR (11).

Nitric oxide (NO), a vasorelaxant has been shown to regulate a variety of physiological functions including platelet aggregation, inflammation, neurotransmission, hormone release, cell differentiation, migration, proliferation and apoptosis (12, 13). Most of the effects have been shown to be mediated through the activation of soluble guanylyl cyclase and cGMP pathways (14); however, other cGMP-independent mechanisms have also been reported (13, 15). VSMC from SHR have been shown to exhibit decreased levels of NO and eNOS (16, 17). We recently showed that augmentation of intracellular levels of NO by a donor, S-Nitroso-N-acetyl-DL-penicillamine (SNAP) attenuated the hyperproliferation of VSMC from SHR by cGMP-independent mechanism (18). We also showed that *in vivo* treatment of SHR with NO donor sodium nitroprusside (SNP) attenuated the development of high blood pressure through the inhibition of overexpression of Giα proteins and nitroxidative stress (19). The present study is therefore undertaken to investigate if *in vivo* treatment of SHR with SNP could also attenuate the hyperproliferation of VSMC and further identify the underlying potential mechanisms involving AT1 receptor, c-Src, growth factor receptors activation, MAP kinase, and cell cycle proteins in mediating this response.

## Materials and Methods

### Chemicals

Sodium nitroprusside (SNP) and losartan were purchased from Sigma-Aldrich Chemical Co. (St Louis, Missouri, USA). Western blotting primary antibodies against cyclin D1 (sc-20,044), cyclin E (sc-481), Cdk2 (sc-6248), Cdk4 (sc-23896), phospho-specific-Ser^249^/Thr^252^ Rb (sc-377528), Rb (sc-102), AT1 (sc-1173), p21 (sc-397), p27 (sc-528), ERK1/2 (sc-135900), p-ERK1/2 (sc-7383, phosphospecific-tyrosine^204^), p-PDGFR-β (sc-12909-R, phosphospecific-tyrosine^1021^), PDGFR-β (sc-432), p-EGFR (sc-101668, phosphospecific-tyrosine^1173^), EGFR (sc-373746), p-IGF-IR (sc-101704, phosphospecific-tyrosine^1165/1166^), IGF-IRβ (sc-713), p-c-Src (sc-166860, phosphospecific-tyrosine^419^), c-Src (sc-18), dynein IC1/2 (sc-13524), secondary antibodies goat anti-mouse IgG horseradish-peroxidase (HRP) conjugate (sc-2005), Donkey anti-goat IgG horseradish-peroxidase (HRP) conjugate (sc-2020) and enhanced chemiluminescence (ECL) detection system kits were purchased from Santa-Cruz Biotechnologies (Santa Cruz, CA, USA). Secondary antibody Goat Anti-Rabbit IgG (H+L)-HRP conjugate was purchased from BIO RAD (USA). The L-(4, 5-^3^H) thymidine was from PerkinElmer Inc. (Waltham, Massachusetts, USA). All other chemicals used in the experiments were purchased from Sigma-Aldrich.

### Animal treatment

Male SHR (8-week-old) and age-matched WKY rats were purchased from Charles River Laboratories International Inc. (St-Constant, Quebec, Canada). Animals were maintained at room temperature with free access to water and regular rat chow in 12 h light/dark cycles. Rats were left for 2 days for acclimatization. SHR and age-matched WKY rats were injected intraperitoneally with SNP (0.5 mg/kg body weight) twice per week for two weeks in 0.01 mol/L sodium phosphate buffer, pH 7.0, containing 0.05 mol/L NaCl. The control WKY rats and SHR received vehicle. The blood pressure (BP) was monitored twice a week by using the CODA noninvasive tail-cuff method, according to the recommendation of American Heart Association (20). At the end of the 10^th^ week, after taking the blood pressure, the rats were euthanized by decapitation after CO_2_ exposure. Thoracic aortas were dissected out and used for cell culture. All the animal procedures used in the present study were approved by the Comité de Déontologie de l’Expérimentation sur les Animaux (CDEA) of the University of Montreal (Approval No. 99050). The investigation conforms to the “Guide for the Care and Use of Laboratory Animals” published by the US National Institutes of Health (NIH) (Guide, NRC 2011).

### Cell culture and incubation

Aortic VSMC from 10 week old control and SNP-treated SHR and their age-matched WKY rats were cultured as described previously (21). The purity of the cells was checked by immunofluorescence technique using α-actin as described previously (22). These cells were found to contain high levels of smooth muscle-specific actin. The cells were plated in 75 cm^2^ flasks and incubated at 37°C in 95% air and 5% CO_2_ humidified atmosphere in Dulbecco’s modified Eagle’s medium (DMEM) (with glucose, L-glutamine and sodium bicarbonate) containing antibiotics and 10% heat-inactivated fetal bovine serum (FBS). The cells were passaged upon reaching confluence with 0.5% trypsin containing 0.2% EDTA and utilized between passages 3 and 10. Confluent cells were then starved by incubation for 4 h in DMEM without FBS at 37°C to reduce the interference by growth factors present in the serum. For the receptor antagonist studies, VSMC from control and SNP-treated SHR and WKY rats were incubated for 24 h in the absence (control) or presence of losartan (10 μM). After incubation, the cells were washed three times with PBS and lysed in 100 µl of buffer (25 mM Tris-HCl, pH 7.5, 25 mM NaCl, 1 mM Na orthovanadate, 10 mM Na fluoride, 10 mM Na pyrophosphate, 2 mM ethylene, bis(oxyethylenenitrolo)tetracetic acid 2 mM ethylenediamine tetracetic acid, 1 mM phenylmethylsulfonyl fluoride, 10 µg/mL aprotinin, 1% Triton X-100, 0.1% sodium dodecyl sulphate (SDS), and 0.5 µg/mL leupeptin) on ice. The cell lysates were centrifuged at 12,000 rpm for 10 min at 4°C, and the supernatants were used for Western blot analysis. Protein concentration was measured with the Bradford assay [23]. Cell viability was checked by the trypan blue exclusion technique as described previously [24] and indicated that >90–95% cells were viable. To examine the role of eNOS in SNP-induced attenuation of hyperproliferation of VSMC from SHR, VSMC from 12 week old male SHR and age-matched WKY rats were preincubated in the absence (control) or presence of Nω-Nitro-L-Arginine Methyl Ester (L-NAME) (100 μM) for 1 h prior to the treatment with SNP (100 μM) for 24 h and were used for cell proliferation study as described below.

### Western blotting

The levels of protein expression and phosphorylation were determined by Western blotting as described previously (23). After SDS-PAGE, the separated proteins were transferred to a nitrocellulose membrane with a semi-dry transblot apparatus (Bio-Rad Laboratories, Mississauga, Ontario, Canada) at 15 V for 45 min or a liquid transfer apparatus (Bio-Rad Laboratories) at 100 V for 1 h. Membranes were blocked for 1 h at room temperature with 5% dry milk and incubated overnight with specific primary antibodies against different proteins in phosphate buffer solution (PBS) containing 0.1% Tween-20 (PBS-T) overnight at 4°C. Dynein was used as loading controls in all experiment. The antibody-antigen complexes were detected by incubating the membranes with horseradish peroxidase-conjugated secondary antibodies for 1 h at room temperature. The blots were then washed three times with PBS-T before reaction with enhanced chemiluminescence (ECL). Quantitative analysis of the proteins was performed by densitometric scanning of the autoradiographs using the enhanced laser densitometer LKB Ultroscan XL and quantified using the gel-scan XL evaluation software (version 2.1) from Pharmacia (Baie d′Urfé, Québec, Canada).

### Determination of cell proliferation

Cell proliferation was quantified by DNA synthesis which was evaluated by incorporation of [^3^H] thymidine into cells as described earlier (23). Subconfluent aortic VSMC from control and SNP-treated SHR and their age-matched WKY rats were plated in 6-well plates for 24 h and were serum deprived for 12 h to induce cell quiescence. The cells were then incubated with [^3^H] thymidine (1 µCi) for 4 h before the cells were harvested. The cells were rinsed twice with ice-cold PBS and incubated with 5% trichloroacetic acid (TCA) for 1 h at 4°C. After being washed twice with ice-cold water, the cells were incubated with 0.4 N sodium hydroxide (NaOH) solution for 30 min at room temperature, and radioactivity was determined by liquid scintillation counter.

### Determination of intracellular levels of nitric oxide (NO)

The levels of intracellular NO produced in VSMC were measured using intracellular fluorescent probes diaminofluorescein-2 diacetate (DAF-2DA) as described earlier (18, 19). Confluent VSMC, after washing twice with PBS, were incubated at 37°C for 1 h with both 10 mmol/L DAF-2DA and 10−6 mol/L acetylcholine for detecting NO. Cells were washed twice with PBS, and fluorescence intensities were measured by a spectrophotometer (TECAN infinite 200 PRO) with excitation and emission wavelengths at 495 nm and 515 nm. Changes in fluorescence intensities were expressed as percentages of the values obtained in the WKY group (taken as 100%).

### Statistical analysis

The number of independent experiments is reported. Each experiment was conducted at least 4 times using separate cell population. All data are expressed as the mean ± SEM. Comparisons between groups were made with a one-way analysis of variance (ANOVA) followed by the Newman-Keuls Multiple Comparison Test, using GraphPad Prism 5 (GraphPad Software Inc., La Jolla, California, USA). Results were considered statistically significant at values of *p*< 0.05.

## Results

The systolic blood pressure (BP) of SHR and WKY rats at 8 weeks were 184.5 ± 10.3 mmHg and 108.5 ± 9.6 mmHg, respectively (*p* < 0.001). Intraperitoneal injection of SNP to 8-week-old SHR (0.5 mg/kg body weight) twice per week decreased the systolic BP in a time-dependent manner and at 10 weeks, the BP was decreased by about 82 mmHg (129.0 ± 5.2 vs. 211 ± 8.8 mmHg, *p* < 0.001) without affecting the BP in WKY rats. In addition, SNP treatment did not have any adverse effects on the health of the animals, because all rats treated with SNP maintained or gained weight during the period of the studies (Body weights at 10 weeks were as follows: WKY rats, 227 ± 5.7 g; SNP-treated WKY rats, 235 ± 3.6 g; SHR, 227 ± 2 g; and SNP-treated SHR, 223 ± 2.4 g). The heart rate was not significantly different between control and SNP-treated groups (Heart rate (Beats\minute) Control WKY rats 364.8 ±4.4, SNP-treated WKY Rats 392.3 ±48.6 : Control SHR,447.9 ±20.5, SNP-treated SHR, 419.6 ±32.4).

### SNP augments the intracellular levels of NO in VSMC from SHR

Our earlier studies have shown that VSMC from SHR exhibit decreased levels of intracellular NO that contribute to the hyperproliferation of VSMC from SHR (18). To investigate if SNP-induced attenuation of VSMC hyperproliferation is due to the augmentation of the intracellular levels of NO, we determined the levels of NO in VSMC from control and SNP-treated SHR and WKY rats. Results shown in Table 1 indicate that the levels of intracellular NO were significantly lower in VSMC from SHR as compared to WKT rats and in vivo treatment of SNP significantly augmented the intracellular levels of NO in VSMC from both WKY and SHR by about 190% and 580%, respectively compared to control groups.

**Table 1:** Effect of *in vivo* sodium nitroprusside (SNP) treatment of SHR on the intracellular levels of nitric oxide (NO) in aortic vascular smooth muscle cells (VSMC) from 10 week old SHR and age-matched WKY rats.

| Groups (N=6) | Mean ± SE (arbitrary units, relative % of WKY group) |
| --- | --- |
| WKY | 100 ± 30.3 |
| WKY+SNP | 293.7 ± 25.2*** |
| SHR | 34.7 ± 8.0** |
| SHR+SNP | 236.8 ± 20.12$\Psi\Psi\Psi$ |

Comparisons between groups were made with a one-way analysis of variance (ANOVA) followed by the Newman-Keuls Multiple Comparison Test,. \*\* $P < 0.01$ , \*\*\* $P < 0.001$ versus WKY; $\Psi\Psi\Psi P < 0.001$ versus SHR.

### SNP attenuates the hyperproliferation of VSMC from SHR

We earlier reported that *in vivo* treatment of SHR with NO donor SNP attenuated the development of high blood pressure (19). Since vascular remodeling due to hyperproliferation is associated with hypertension, it was of interest to investigate if *in vivo* treatment of SHR with SNP could also attenuate the hyperproliferation of VSMC from SHR. As shown in Figure 1, the proliferation of VSMC from SHR was significantly augmented by about 100% as compared to WKY rats as determined by [^3^H] thymidine incorporation, and SNP treatment attenuated it by about 50%. On the other hand, SNP did not have any effect on the proliferation of VSMC from WKY rats.

**Figure 1:**
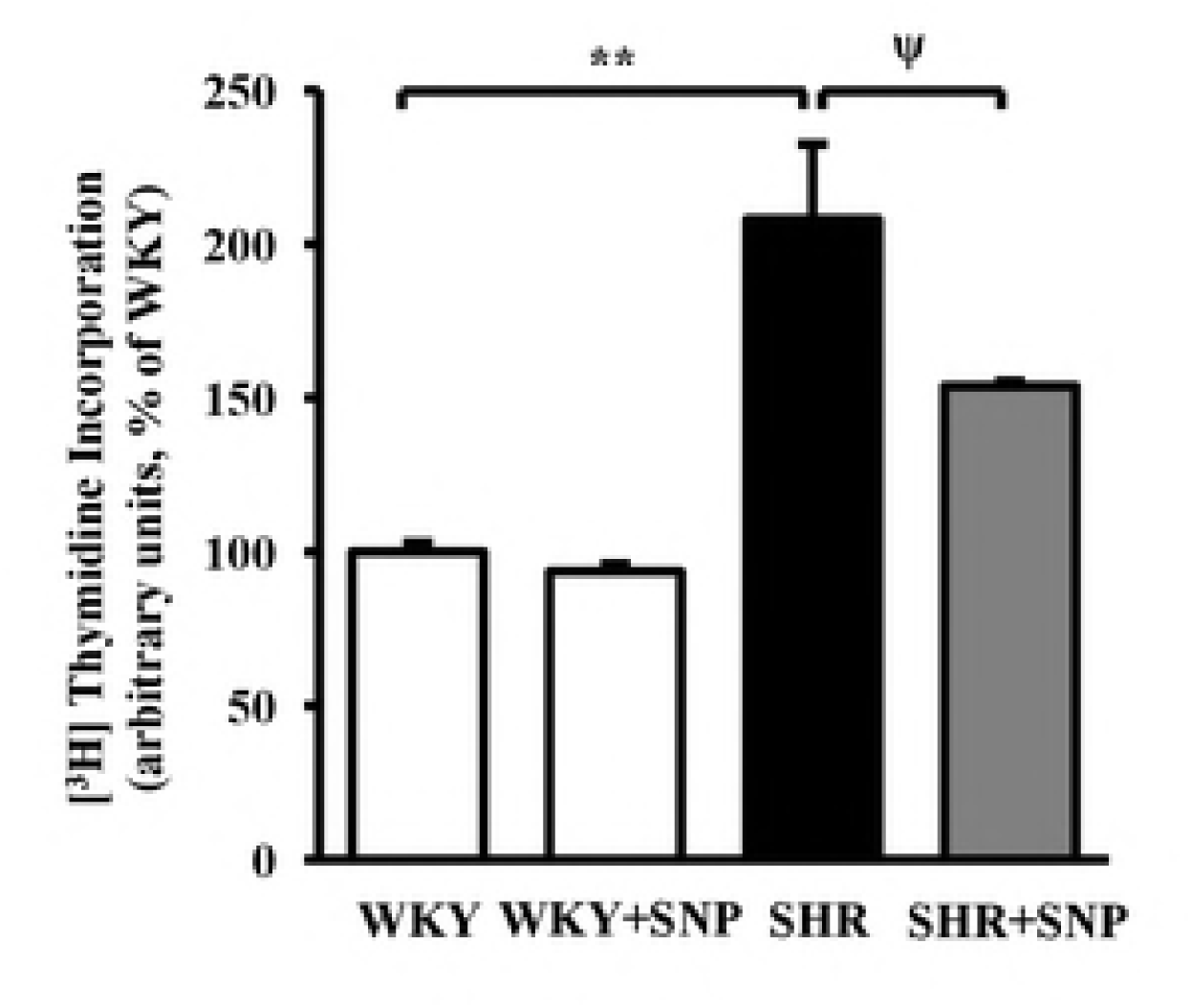
Effect of *in vivo* SNP treatment on DNA synthesis in aortic VSMC from SHR and age-matched WKY rats. Aortic VSMC from 10 week old SHR and age-matched WKY rats with or without SNP-treated groups were cultured and [^3^H] thymidine incorporation was determined as described in the “Materials and Methods” section. Values are mean ± SEM of 6 independent experiments. The results are expressed as percentage of WKY (control) which has been taken as 100%. \*\**p*<0.01 versus WKY, ^ψ^*p*<0.05 versus SHR.

### Implication of eNOS in SNP-induced antiproliferative effect in VSMC from SHR

We earlier showed that *in vivo* treatment of SHR with SNP that elevated the intracellular levels of NO also augmented the reduced levels of intracellular eNOS in VSMC from SHR (19). To investigate the contribution of SNP-induced enhanced levels of intracellular eNOS in the antiproliferative effect of SNP, we examined the effect of pretreatment of VSMC from SHR and WKY rats with L-NAME, a non-selective inhibitor of NOS (24) on the antiproliferative effect of SNP and the results are shown in Figure 2. As reported earlier (18), SNP inhibited the hyperproliferation of VSMC from SHR by about 40%, however, the inhibition of NOS by pretreatment with L-NAME, augmented the hyperproliferation in VSMC from SHR and reversed the antiproliferative effect of SNP towards SHR (control) levels. On the other hand, these treatments did not have any significant effect in VSMC from WKY rats.

### SNP attenuates the expression of cell cycle proteins in VSMC from SHR

The enhanced expression of cyclin D1, cyclin E and Cdk2 was shown to contribute to the hyperproliferation of VSMC from SHR (10). To investigate if SNP-induced attenuation of hyperproliferation of VSMC from SHR is attributed to the inhibition of the enhanced expression of cell cycle proteins, we examined the effect of SNP on the levels of different cell cycle proteins. As shown in Figure 3, the expression of cyclin D1 (A), cyclin E (B) Cdk2 (C) and Cdk4 (D) was significantly enhanced by about 110%, 115%, 100% and 60% respectively in VSMC from SHR as compared to WKY and SNP attenuated the enhanced expression of cyclin D1 and Cdk4 to control levels whereas the increased expression of cyclin E and Cdk2 was inhibited by about 80% and 50% respectively. In addition, SNP also decreased the expression of cyclin D1 in VSMC from WKY rats by about 40%. Furthermore, the expression of phosphorylated retinoblastoma protein (pRB) (E) that was enhanced by about 180% in VSMC from SHR was also decreased by SNP treatment by about 55%, however, SNP did not affect the levels of pRB in WKY rats. In addition, the levels of RB were not different in VSMC from SHR from WKY rats and SNP did not have any effect on the expression of RB proteins in VSMC from both SHR and WKY rats (E).

Furthermore, the levels of Cdk inhibitors, p21^Cip1^ (Figure 4A) and p27^Kip1^ (Figure 4B) were significantly decreased by about 65% and 35% respectively in VSMC from SHR as compared to WKY and SNP treatment restored the reduced expression of these inhibitors by about 20% and 80% respectively. On the other hand, SNP treatment did not have any significant effect on the levels of these proteins in WKY rats.

### SNP attenuates the enhanced expression of AT1 receptor in VSMC from SHR

Since enhanced levels of endogenous Ang II through the activation of AT1 receptors were shown to contribute to the hyperproliferation of VSMC from SHR (25), it was of interest to examine if SNP-induced inhibition of hyperproliferation of VSMC from SHR is attributed to its ability to attenuate the enhanced expression of AT1 receptor and AT1 receptor-induced enhanced expression of cell cycle proteins. To test this, we examined the expression of AT1 receptor in VSMC from control and SNP-treated SHR and WKY rats. Results shown in Figure 5A, indicate that the expression of AT1 receptor is significantly augmented by approximately 400% in VSMC from SHR as compared to WKY rats and SNP treatment restored the enhanced expression of AT1 receptor to WKY control level whereas the expression of AT1 receptor in VSMC from WKY rats was not affected by SNP treatment

### Role of AT1 receptor in enhanced expression of cell cycle proteins

To further investigate the contribution of enhanced expression of AT1 receptor in enhanced expression of cell cycle proteins in VSMC from SHR, we examined the effect of losartan, an AT1 receptor antagonist on the expression of cell cycle proteins in VSMC from SHR and WKY rats. Results shown in Figure 5B indicate that treatment of VSMC from SHR with losartan significantly attenuated the overexpression of cyclin D1 by about 60% whereas the enhanced expression of Cdk2 (5C) and Cdk4 (5D) was completely restored to control levels. On the other hand, losartan did not have any effect on the expression of these proteins in VSMC from WKY rats. These results suggest the involvement of AT1 receptor in the overexpression of cell cycle proteins in VSMC from SHR.

### SNP attenuates the enhanced activation of c-Src in VSMC from SHR

Since the activation of AT1 receptor through oxidative stress activates c-Src in VSMC from SHR and c-Src is implicated in the overexpression of cell cycle proteins (11) and hyperproliferation (7), it was of interest to examine if SNP inhibits hyperproliferation through the inhibition of c-Src activation in VSMC from SHR. Results shown in Figure 6 indicate that the phosphorylation of Tyr^419^ on c-Src was significantly augmented by about 70% in VSMC from SHR compared with WKY rats and SNP attenuated the enhanced phosphorylation of c-Src by about 65%. In addition, SNP also decreased the phosphorylation of c-Src in VSMC from WKY rats by approximately 20%.

### SNP attenuates the enhanced activation of growth factor receptors in VSMC from SHR

Earlier studies showed the contribution of growth factor receptor, downstream molecules of c-Src in the enhanced expression of cell cycle proteins (11) and hyperproliferation of VSMC from SHR (6, 7). To further examine if SNP-induced attenuation of hyperproliferation of VSMC from SHR is also mediated through the inhibition of EGF-R, PDGF-R, and IGF-IR activation, we investigated the effect of SNP on the phosphorylation of EGF-R, PDGF-R, and IGF-IR in VSMC from SHR and WKY rats using specific phosphotyrosine antibodies. As shown in Figure 7, phosphospecific-Tyr^1173^-EGF-R (A), phosphospecific-Tyr^1021^-PDGF-R (B) and phosphospecific-Tyr^1165/1166^-IGF-IR (C) detected a single band of 150, 190 and 97 kDa corresponding to EGF-R, PDGF-R and IGF-IR, respectively, in VSMC from both SHR and WKY rats. However, the extent of EGF-R, PDGF-R and IGF-IR phosphorylation was greater by about 430, 130, and 90%, respectively, in VSMC from SHR compared with VSMC from WKY rats and SNP treatment attenuated the increased phosphorylation of EGF-R, PDGF-R, and IGF-IR to WKY control levels. However, this treatment did not have any significant effect on the phosphorylation of EGF-R, PDGF-R and IGF-IR in VSMC from WKY rats. In addition, the expression of total EGF-R, PDGF-R and IGF-IR was not altered in VSMC from SHR compared with WKY.

### SNP attenuates the enhanced activation of ERK1/2 in VSMC from SHR

The implication of enhanced phosphorylation of ERK1/2 in hyperproliferation (5) and overexpression of cell cycle proteins in VSMC from SHR (10) has been reported. In addition, NO was shown to regulate the activity of MAP kinase in VSMC from SHR (26-28). To investigate if SNP-evoked attenuation of hyperproliferation is due to the inhibition of MAP kinase activation in VSMC from SHR, we examined the effect of SNP on ERK1/2 phosphorylation in VSMC from SHR and the results are shown in Figure 8. As reported earlier (5, 26, 29), the phosphorylation of ERK1/2 was significantly augmented by about 230% in VSMC from SHR compared with WKY and SNP treatment attenuated the enhanced phosphorylation of ERK1/2 by about 65%. However, this treatment did not have any significant effect on ERK1/2 phosphorylation in VSMC from WKY rats.

## Discussion

We and others reported earlier that VSMC from SHR exhibit exaggerated cell growth (proliferation) compared to VSMC from WKY rats (4, 5, 25) which was shown to be attributed to the decreased levels of endogenous NO because elevating the intracellular levels of NO by NO donor, SNAP, attenuated the hyperproliferation by cGMP-independent mechanism (18). However, in the present study we show for the first time that *in vivo* treatment of SHR with SNP that was shown to decrease the high blood pressure (19) also attenuates the hyperproliferation of VSMC by inhibiting the overexpression of AT1 receptor, cell cycle proteins, c-Src, EGF-R, PDGF-R and IGF-IR transactivation and MAP kinase signaling.

Several *in vitro* studies have demonstrated that NO and NO donors inhibit proliferation of various cell types including VSMC by cGMP-independent mechanism (18, 30-32). SNP was shown to attenuate the hyperproliferation of aortic VSMC induced by EGF (33). In addition, Etienne et al. have also demonstrated the antiproliferative effect of SNP in aortic VSMC from diabetic rats (34). However, we report for the first time that *in vivo* treatment of SHR with SNP that elevated the intracellular levels of NO also attenuates the hyperproliferation of aortic VSMC. The fact that the antiproliferative effect of SNP in VSMC from SHR was reversed by the inhibition of NOS by L-NAME further suggests the implication of NOS in SNP-induced elevated levels of intracellular NO and antiproliferation of VSMC. In support of this, *in vivo* treatment of SHR with SNP has been shown to augment the reduced levels of eNOS in VSMC (19). In addition, we also demonstrate that the antiproliferative effect of SNP is associated with the inhibition of overexpression of cell cycle proteins including cyclin D1, cyclin E, Cdk2, Cdk4 and pRB and augmentation of the downregulated levels of Cdk inhibitors p21^Cip1^ and p27^Kip1^ from G1 to S phase. In this regard, the implication of cell cycle proteins from G1-S phase in cell proliferation is well established (35). Furthermore, Ang II- and FBS-induced exaggerated growth of VSMC from SHR was also shown to be associated with progression from G1 to S phase (9, 36). We recently showed the contribution of overexpression of cell cycle proteins from G1-S phase in hyperproliferation of VSMC from SHR (10, 11). Taken together, it may be suggested that the antiproliferative effect of SNP in VSMC from SHR may be attributed to its ability to attenuate the overexpression of cell cycle proteins and upregulation of Cdk inhibitors p21^Cip1^ and p27^Kip1^ from G1 to S phase. In this regard, the role of downregulation of Cdk2 activity and cyclin A gene transcription in antiproliferative effect of NO in VSMC has also been reported (37). In addition, Ishida el al. (38) has demonstrated that the antiproliferative effect of NO donor, SNAP, was associated with the induction of Cdk inhibitor p21^Sdi1/Cip1/Waf1^ in VSMC (38).

The implication of vasoactive peptides in the proliferation of VSMC has been well documented (23, 39). We earlier reported that Ang II, ET-1 and arginine-vasopressin (AVP) increased the proliferation of A10 VSMC via Giα/MAP kinase pathways (23). In addition, Ang II treatment of VSMC from SHR was shown to enhance the proliferation to a greater extent compared to VSMC from WKY rats which was attenuated by Ang II AT1 receptor antagonist, losartan, indicating the implication of AT1 receptor in the enhanced proliferation of VSMC from SHR (40). Furthermore, the enhanced levels of endogenous Ang II through the activation of AT1 receptor and associated signaling have also been shown to contribute to the hyperproliferation of VSMC from SHR (25). The fact that *in vivo* treatment of SHR with SNP attenuates the overexpression of AT1 receptor and hyperproliferation of VSMC further suggests that SNP-induced attenuation of hyperproliferation may be attributed to its ability to downregulate the enhanced expression of AT1 receptor. These results are consistent with our earlier *in vitro* study showing that treatment of VSMC from SHR with SNAP, another donor of NO for 24 h attenuated the overexpression of AT1 receptor and hyperproliferation (18). In addition, NO-mediated inhibition of Ang II binding (41) and AT1 receptor-induced migration of aortic VSMC has also been reported (42). In the present study, we also demonstrate the implication of AT1 receptor in enhanced expression of cell cycle proteins in VSMC from SHR because losartan, AT1 receptor antagonist attenuates the overexpression of cell cycle proteins. Taken together, it may be suggested that SNP-induced antiproliferative effect is mediated through the downregulation of AT1 receptor and attenuation of enhanced expression of cell cycle proteins.

Earlier studies have demonstrated that the enhanced levels of endogenous Ang II through the interaction with AT1 receptor increase oxidative stress which via the activation of c-Src, growth factor receptor and MAP kinase signaling pathways increase the expression of Giα proteins and result in the hyperproliferation of VSMC from SHR (25). In addition, the role of oxidative stress, c-Src, growth factor receptors, MAP kinase and Giα proteins in the overexpression of cell cycle proteins in VSMC from SHR has also been demonstrated (10, 11). Here we show for the first time that *in vivo* treatment of SHR with SNP inhibits the enhanced activation (phosphorylation) of c-Src, growth factor receptors EGF-R, PDGF-R, and IGF-IR as well as ERK1/2, all the signaling molecules implicated in the proliferation and expression of cell cycle proteins from G1-S phase (10, 11, 43). These results suggest that the inhibition of enhanced activation of c-Src, growth factor receptor activation and MAP kinase by SNP attenuates the overexpression of cell cycle proteins in VSMC from SHR and result in the attenuation of hyperproliferation.

## Conclusions

In conclusion, we provide the evidence that in vivo treatment of SHR with SNP attenuates the enhanced expression of AT1 receptor, c-Src and growth factor receptor activation as well as MAP kinase signaling, all these signaling molecules through the inhibition of overexpression of cell cycle proteins result in the suppression of hyperproliferation of VSMC. Thus, it can be suggested that NO donors including SNP by elevating the intracellular levels of NO may contribute to the amelioration of vascular remodelling and may thus be used as potential therapeutic agents for the treatment of vascular complications of hypertension.

## Acknowledgements

We also thank the Fonds de recherche du Québec-Nature et technologies (FRQNT) for the merit scholarship (PBEEE) to Ekhtear Hossain at post-doctoral (V2) category (File No.200683).

---

**Figure 2: Effect of L-NAME on SNP-induced attenuation of DNA synthesis in aortic VSMC from SHR and age-matched WKY rats.** Aortic VSMC from 12 week old SHR and age-matched WKY rats after pretreatment with L-NAME (100μM) for 1 h were exposed to SNP (100 μM) for 24 h.and [3H] thymidine incorporation in these cells was determined as described in the “Materials and Methods” section. Values are mean ± SEM of 6 independent experiments. The results are expressed as percentage of WKY (control) which has been taken as 100%. **p<0.01, ***p <0.001.

**Figure 3: Effect of *in vivo* SNP treatment on cell cycle proteins: Cyclin D1, Cyclin E, Cdk2, Cdk4, and pRB in aortic VSMC from SHR and age-matched WKY rats.** Aortic VSMC lysates from 10-week-old SHR and WKY rats with or without SNP treatment were subjected to Western blot analysis using specific antibodies against Cyclin D1 (A), Cyclin E (B), Cdk2 (C), Cdk4 (D) and phosphorylated pRB/total RB (E) as described in the “Materials and Methods” section. Dynein was used as the loading control. Lower panels show the quantification of protein bands by densitometric scanning. Values are mean ± SD of 4 independent experiments. The results are expressed as percentage of WKY (control) which has been taken as 100%. \**p*<0.05, \*\**p*<0.01, \*\*\**p*<0.001 versus WKY; ^ψ^*p*<0.05, ^ψψ^*p*<0.01, ^ψψψ^*p*<0.001 versus SHR.

**Figure 4: Effect of *in vivo* SNP treatment on Cdk inhibitors: p21^Cip1^ and p27^Kip1^ in aortic VSMC from SHR and age-matched WKY rats.** Aortic VSMC lysates from 10-week-old SHR and WKY rats with or without SNP treatment were subjected to Western blot analysis using specific antibodies against p21^Cip1^ (A) and p27^Kip1^ (B) as described in the “Materials and Methods” section. Dynein was used as the loading control. Lower panel shows the quantification of protein bands by densitometric scanning. Values are mean ± SD of 4 independent experiments. The results are expressed as percentage of WKY (control) which has been taken as 100%. \**p*<0.05, \*\*\**p*<0.001 versus WKY; ^ψ^*p*<0.05 versus SHR.

**Figure 5: (A) Effect of *in vivo* SNP treatment on the expression of AT1 receptor in aortic VSMC from SHR and age-matched WKY rats.** Aortic VSMC lysates from 10-week-old SHR and WKY rats with or without SNP treatment were subjected to Western blot analysis using specific antibodies against AT1R as described in the “Materials and Methods” section. Dynein was used as the loading control. Lower panels show the quantification of protein bands by densitometric scanning. **(B-D) Effect of losartan on the expression of cell cycle proteins: Cyclin D1, Cdk2, Cdk4 in aortic VSMC from SHR and age-matched WKY rats.** Confluent aortic VSMC from 10-week-old SHR and WKY rats were starved for 4 h and incubated in the absence (control) or presence of losartan (LOS, 10 µM) for 24 h. The cell lysates were prepared and subjected to Western blot analysis using specific antibodies against Cyclin D1 (B), Cdk2 (C), and Cdk4 (D) as described in the" Materials and Methods" section. Values are mean ± SD of 4 independent experiments. The results are expressed as percentage of WKY (control) which has been taken as 100%. **p<0.01, ***p<0.001 versus WKY; ^ψψψ^ p<0.001 versus SHR.

**Figure 6: Effect of *in vivo* SNP treatment on the phosphorylation of c-Src in aortic VSMC from SHR and age-matched WKY rats.** Aortic VSMC lysates from 10-week-old SHR and WKY rats with or without SNP treatment were subjected to Western blot analysis using specific antibodies against phosphorylated c-Src and total c-Src as described in the “Materials and Methods” section. Lower panel shows the quantification of protein bands by densitometric scanning. Values are mean ± SD of 4 independent experiments. The results are expressed as percentage of WKY (control) which has been taken as 100%. \*\**p*<0.01 versus WKY, ^ψ^*p*<0.05 versus SHR.

**Figure 7: Effect of *in vivo* SNP treatment on the phosphorylation of EGF-R, PDGF-R, and IGF-IR in aortic VSMC from SHR and age-matched WKY rats.** Aortic VSMC lysates from 10-week-old SHR and WKY rats with or without SNP treatment were subjected to Western blot analysis using specific antibodies against phosphorylated EGF-R/total EGF-R (A), phosphorylated PDGF-R/total PDGF-R (B) and phosphorylated IGF-IR/total IGF-IR (C) as described in the “Materials and Methods” section. Lower panels show the quantification of protein bands by densitometric scanning. Values are mean ± SD of 4 independent experiments. The results are expressed as percentage of WKY (control) which has been taken as 100%. \*\**p*<0.01 \*\*\**p*<0.001 versus WKY; ^ψ^*p*<0.05, ^ψψ^*p*<0.01 versus SHR.

**Figure 8: Effect of *in vivo* SNP treatment on the phosphorylation of ERK 1/2 in aortic VSMC from SHR and age-matched WKY rats.** Aortic VSMC lysates from 10-week-old SHR and WKY rats with or without SNP treatment were subjected to Western blot analysis using specific antibodies against phosphorylated ERK1/2 and total ERK1/2 as described in the “Materials and Methods” section. Lower panel shows the quantification of protein bands by densitometric scanning. Values are mean ± SD of 4 independent experiments. The results are expressed as percentage of WKY (control) which has been taken as 100%. \*\**p*<0.01 versus WKY, ^ψψ^*p*<0.01 versus SHR.

## Abbreviations

SNP: sodium nitroprusside
ODQ: H (1, 2, 4) oxadiazole (4, 3-a) quinoxalin-1-one
Ang II: angiotensin II
AT1 receptor: angiotensin II type 1 receptor
ET-1: endothelin-1
WKY rat: Wistar-Kyoto rat
SHR: spontaneously hypertensive rats
EGFR: epidermal growth factor receptor
PDGFR: platelet-derived growth factor receptor
IGF-IR: insulin-like growth factor 1 (IGF-1) receptor

